# Full length TECPR1 displays ‘cis’ Dysferlin domain architecture

**DOI:** 10.64898/2026.03.13.711659

**Authors:** Ernest A. Okertchiri, John B. Miles, C. Keith Cassidy, Adam L. Yokom

**Author notes:** Correspondence: Adam Lee Yokom.

## Abstract

Tectonin Beta-Propeller Repeat containing 1 (TECPR1) is an essential regulator of a noncanonical autophagy pathway known as Sphingomyelin TECPR1 induced LC3 lipidation (STIL). TECPR1 forms an E3-like ligase complex and recognizes exposed sphingomyelin on damaged membranes. TECPR1 contains five folded domains however the structural basis for TECPR1 function has remained unresolved. Here, we report the first structure of full length TECPR1 resolved using cryo electron microscopy. TECPR1 forms an elongated hook shaped architecture that positions Dysferlin domains in a cis arrangement. Our structure uncovers an uncharacterized intramolecular interface between tectonin repeat 1 and PH domains. This interaction forms a stabilizing bridge that contributes to the orientation of the DysF domains. Molecular dynamics simulations further demonstrate that TECPR1 maintains the overall structural arrangement during membrane association. Our data provide a structural framework for how TECPR1 domain arrangement corresponds with membrane binding.

## Introduction

Autophagy (also termed macroautophagy) is an essential cellular process which functions to maintain cellular homeostasis via degradation of intercellular debris. Autophagy encompasses a core set of autophagy-related proteins which are necessary for initiation, elongation, closure, and fusion along with auxiliary adaptor and effector proteins. A hallmark for autophagy is the conjugation of ubiquitin-like ATG8 family proteins (LC3s and GABARAPs in mammals) to lipid headgroups ^1^. ATG8 homologs are attached to phosphatidylethanolamine or phosphatidylserine, marking the autophagosome membrane as it is built de novo. In most autophagy modalities, ATG16L1 forms a complex with ATG5 and ATG12 to accomplish this crucial conjugation step ^2^.

Recent work has highlighted non-canonical autophagy-pathways that adapt core autophagy machinery to more specialized ATG8 conjugation pathways ^3^. Conjugation of ATG8s to Single Membranes (CASM) is a general term for pathways which function independent from traditional autophagy signaling cascades ^4,5^ Unlike macroautophagy, in which ATG8 proteins are conjugated to the double-membraned phagophore, CASM involves the lipidation of ATG8s onto pre-existing single-membrane compartments. CASM is activated under many forms of cellular stress. This includes damage to acidic compartments where resident V-ATPase complexes recruit the ATG16L1:ATG5-ATG12 E3-like complex ^6^. The role of Tectonin Beta-Propeller Repeat-containing 1 (TECPR1) as a key protein of Sphingomyelin-TECPR1-induced LC3 lipidation (STIL) has been recently established ^7–9^ As the name implies, STIL occurs during the exposure of sphingomyelin to the cytosol, such as when the lysosomal membrane ruptures during bacterial infection. TECPR1 was initially identified as an autophagy-related protein based on an interaction with ATG5 ^10^. Subsequent studies revealed that TECPR1 can replace ATG16L1 in the E3-like ligase complex and promote LC3 conjugation independently ^11,12^ TECPR1 conjugation occurs during membrane damage and is estimated to account for ∼10-20% of LC3 conjugation when ATG16L1 is present ^7,8,13^.

TECPR1 is a large multi-domain protein composed of six structural domains, two Dysferlin (DysF) domains, two tectonin repeat domains, an ATG5-interacting region (AIR) and a plekstrin homology (PH) domain (Figure 1A) ^14–16^. DysF domains are associated with membrane binding and repair and are found across myoferlin and Ferlin proteins ^17,18^. TECPR1 DysF domains have been identified as the sites of sphingomyelin (SM) binding and thus are responsible for recruitment of TECPR1 to the membrane surface. Each dysferlin domain contains a hydrophobic tip with aromatic amino acids (W77, W154 in DysF1 and W829, F908 in DysF2) responsible for SM binding ^7,13^. The mechanism of TECPR1 membrane recruitment remains debated. The DysF1 domain has been reported to be sufficient for SM-dependent recruitment with minimal contribution from the DysF2 domain, supporting a trans-like membrane engagement ^7^. In contrast, both DysF domains cooperate in membrane binding, which is consistent with a cis-like model ^9,13^. These findings highlight an unresolved question on whether TECPR1 engages damaged membranes through a single DysF domain or coordinated action of both domains. TECPR1 contains two WD40-repeat domains (TR1 and TR2). Proteins containing WD40 domains have been described to mediate interactions with other proteins, peptides and are involved in signal transduction processes ^19,20^. The N-terminal TR1 domain possesses an LC3C binding site that targets TECPR1 to lysosomes while the C-terminal TR2 domain is localized in the cytosol ^21^. In addition to membrane recruitment, TECPR1 contains an ATG5-interacting region (AIR) that allows TECPR1 to recruit the ATG5-ATG12 conjugate directly. Crystallographic structures determined that TECPR1 AIR binds ATG5-ATG12 in the same manner as ATG16L1 ^22^. Finally, TECPR1 encodes a PH domain which is suggested to bind phosphatidylinositol 3-phosphate (PI3P). This was observed only in the presence of the ATG5-ATG12 conjugate ^15^. Despite the established role of TECPR1 in STIL, the structural architecture and in turn regulation of LC3 conjugation remains poorly described. Even in the age of artificial intelligence assisted protein predictions, it remains unclear how the dysferlin domains are oriented in the full length TECPR1 structure or how this positioning might contribute to membrane recruitment.

**Figure 1.**
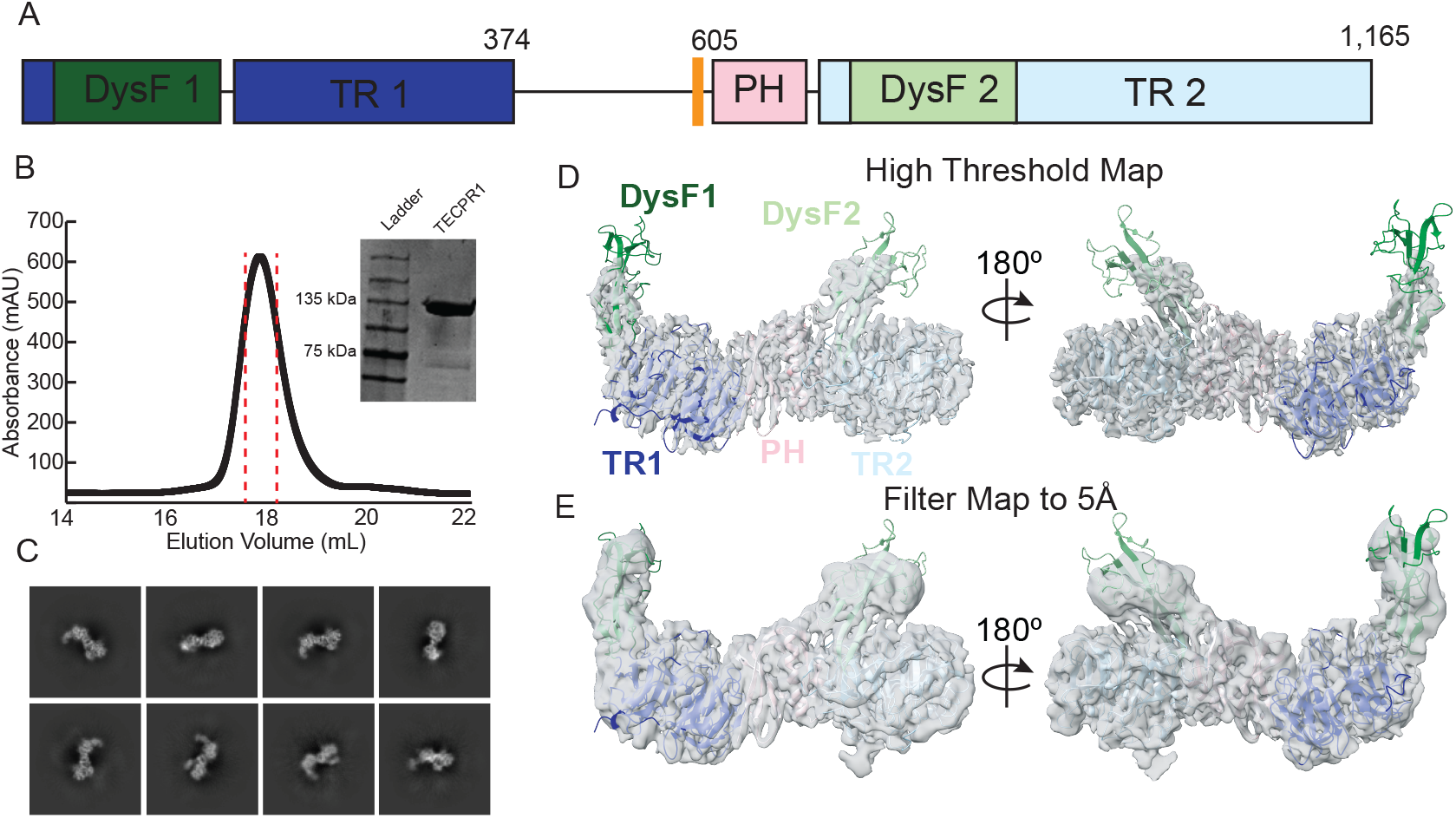
Structural characterization of human TECPR1. **A**) Domain architecture of human TECPR1 colored according to domain (TR1-dark blue, DysF1 – dark green, AIR helix – orange, PH – pink, TR2 – light blue, DysF2 – light green). Residue boundaries are indicated. **B**) Size-exclusion chromatography profile of strep tagged TECPR1. Fractions pooled for further studies are shown with red dashed lines. Inset of SDS–PAGE gel analysis of the peak fraction which shows a molecular weight corresponding to full length TECPR1, ∼135 kDa. **C**) Representative 2D class averages for the combined zero degree and tilted (30°) datasets. **D**) CryoEM density map of TECPR1 displayed at high threshold, with structure coordinates colored as in the domain schematic, Figure 1a. **E**) A 5 Å filtered density map is shown in corresponding orientations with low resolution density present for all domains (DysF1, TR1, PH, DysF2, TR2).

In this study, we have utilized cryo electron microscopy (CryoEM) to determine the first full length structure of human TECPR1. Overall, TECPR1 forms an elongated hook shape bridged by a novel interface between the TR1 and the PH domains. This arrangement positions the DysF domains in a ‘cis’ conformation which are primed for membrane recruitment. Furthermore, all atom simulations suggest that DysF remain docked to the membrane simultaneously and that the TR1-PH interaction is maintained during membrane binding. Together, these data support the idea that TECPR1 functions via multiple lipid interactions at the membrane surface, and that positioning of the DysF domains is regulated by the intramolecular TR1-PH interaction.

### Results– TECPR1 forms elongated hook architecture

Full length TECPR1 was transiently expressed in suspension HEK293F cells and isolated using Strep-Tactin affinity purification. TECPR1 was robustly produced and TECPR1 was further purified using Size Exclusion Chromatography (Figure 1B). SDS-PAGE gel analysis showed a singular band confirming successful isolation of homogeneous human TECPR1. TECPR1 was taken directly to vitrification for single particle CryoEM analysis. Initial screening attempts revealed that TECPR1 had a severe orientation bias, and 3D reconstructions were unsuccessful. Subsequent data collection was performed with a stage tilt of 30 degrees along with the addition of 0.01 mM Lauryl Maltose Neopentyl Glycol. Both modifications mitigated the preferred orientation and increased the number of single particles. Iterative 2D classification revealed a distinct hook shape along with elongated density (Figure 1C, Figure S1A). From these data, a final reconstruction at a nominal resolution of 3.26 Å was determined (Figure 1D, Figure S1B, Table S1). Notably, some orientation bias remained, and the local resolution varies heavily across the map (from 3.5 Å to 12 Å) (Figure S1C-D). The ‘hook’ region seen in 2D and 3D data corresponded with the DysF1 domain (Figure 1E, Movie S1). Additionally, DysF2 is observed in a ‘cis’ position relative to DysF1 positioning both DysF domains along the same face of a TECPR1 structure. We suspect this spatial organization may permit a cooperative membrane engagement similar to how scaffold proteins involved in ATG8 lipidation arrange lipid binding domains and elements to enhance avidity ^23,24^. At a local resolution level (Figure S1C), DysF domains displayed lower resolution and beta strands where not distinguishable when filtered to high resolution. Given that hydrophobic domains are known to interact with the air water interface, we suspect that the DysF domains drive the preferred orientation of TECPR1 ^25^. However, when filtering our cryoEM density to 5 Å, the DysF domains are seen as two distinct densities (Figure 1E). The absence of density corresponding to the AIR helix further indicates the dynamic nature of the linker between the AIR and PH domain (Figure 1A), raising the possibility that this region may become structured upon interaction with the ATG5-ATG12 conjugate. Collectively, this structure shows that TECPR1 exists in an extended conformation in the absence of autophagy binding partners.

### TR1-PH interface form intraprotein bridge

The elongated architecture of TECPR1 is rooted in a novel intramolecular interface between the tectonin repeat 1, TR1 and PH domains. This interaction forms a structural bridge linking the two folded halves of TECPR1 and contributes to the alignment of the DysF domains (Figure 2A). The TR1-PH interface is formed by two loops located between blades 5 and 6 of the first tectonin repeat domain. These loop regions engage the PH domain by blocking its predicted lipid-binding pocket. Multisequence alignment of vertebrate TECPR1 sequences (Figure S2A) reveals strong conservation within the PH domain of TECPR1 (residues 611-717). However, a high sequence conservation is observed among PH domains ^26,27^. Intriguingly, residues within the TR1 loop regions and the adjacent β-barrel surface of the PH domain are likewise conserved (Figure S2A). This conservation suggests functional importance and that the TR1-PH interface is not an artifact but rather is essential for TECPR1’s cellular function ^9,15^. The TR1-PH bridge is composed of a mix of hydrophobic and electrostatic residues that provide 712 Å^2^ of buried surface area (Figure 2A). At the center of the TR1-PH bridge lies a hydrophobic core (Figure 2A) formed by residues V610, F648, and Y650 within the PH domain, positioned on the back side of the putative lipid binding site, together with I329 and L367 located in the TR1 loops. These residues are conserved across the limited number of sequences available for TECPR1 (Figure S2A). This central hydrophobic patch is flanked by polar and electrostatic contacts from the TR1 domain surface (N319, N322, H321 and S368) (Figure 2A). Beyond structural stabilization, the TR1-PH interface may represent an autoinhibitory state that limits PH domain accessibility until conformation rearrangement is triggered. Previous studies have revealed that the interaction of ATG5-ATG12 might expose the PH domain of TECPR1 to bind PI(3)P and PI(4)P directly ^14,15^. In this model, disruption of this TR1-PH interface could be the trigger that couples membrane recognition to downstream ATG8 lipidation activity. Therefore, the TR1-PH bridge may function not only as structural support but also to regulate downstream LC3 conjugation.

**Figure 2.**
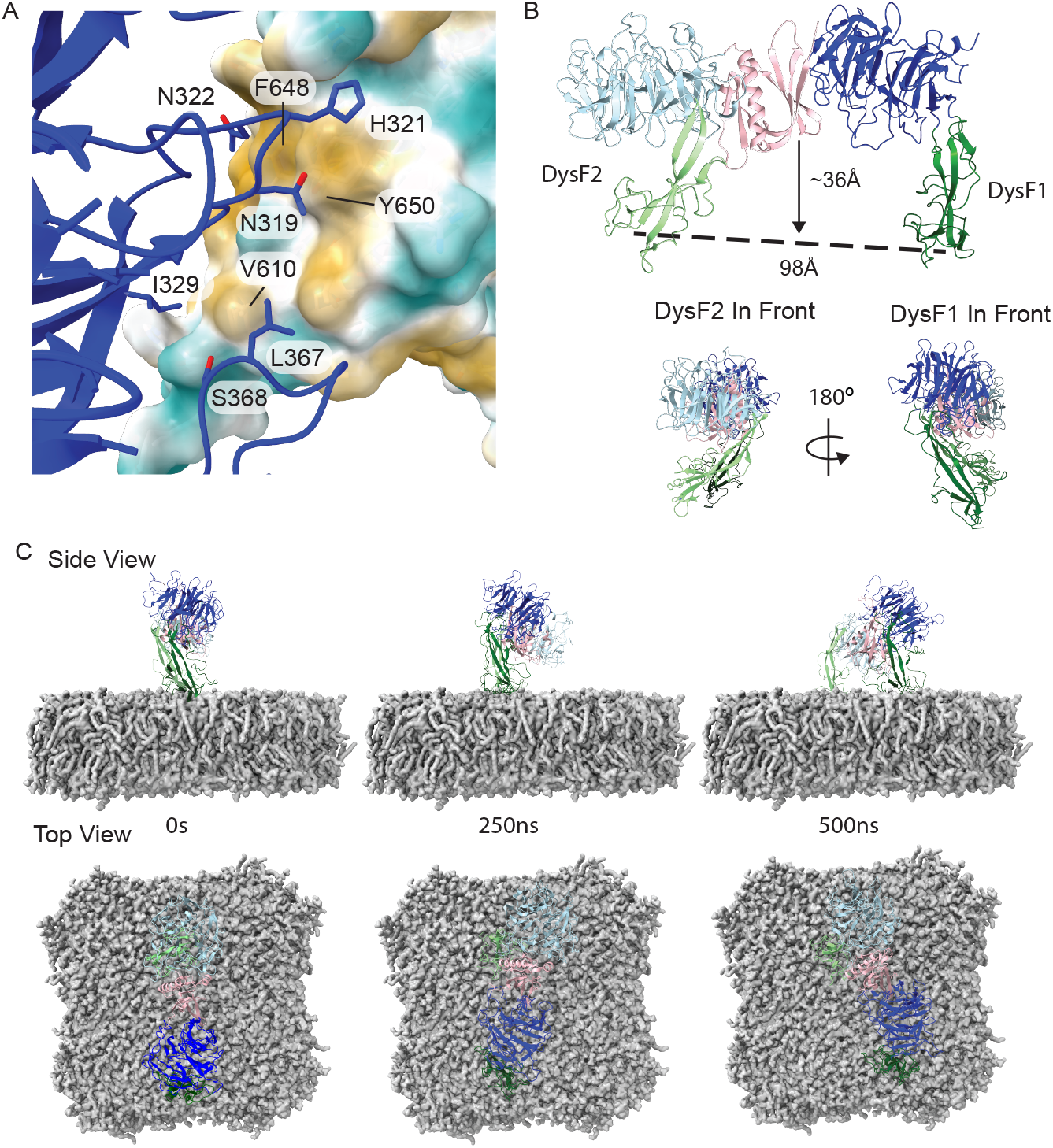
TECPR1 structure reveals TR1-PH bridge and cis Dysferlin domains. **A**) Close-up view of the intra-domain interface between TR1 and PH (shown with hydrophobic surface). Key residues contributing to the interface are labeled, including N319, N322, I329, H321, L367, S368 (TR1 region) and V610, F648, Y650 (PH region). **B**) The relative positioning of the two DysF domains is shown in both front and side views. The two DysF domains align within the same plane and have sphingomyelin binding sites positioned 98 Å apart. The putative lipid binding site in the PH domain is ∼36 Å away from this lipid binding plane between DysF domains. **C**) Molecular dynamics simulation snapshots of human on a lipid surface (7 POPC:2 POPE:1 DPSM lipid ratio) shown at 0 ns, 250 ns, and 500 ns. The overall architecture remains stable relative to the cryo EM structure. There is a progressive diffusion of DysF1 domain across the membrane surface.

### Dysferlin domains form “cis” membrane binding surface

An immediate observation of the full length TECPR1 structure is the spatial arrangement of the DysF domains (Figure 1D). Both DysF1 and DysF2 domains protrude from the central core of TECPR1 and aligned along the same molecular face (Figure 2B). DysF1 and DysF2 are pointed away from their respective TR domains. This places the DysF domains in a position that is compatible with simultaneously membrane engagement. The DysF domains displayed lower resolution (7 Å -12 Å) when compared to the core of TECPR1 reconstruction (Figure S1C) suggesting conformational flexibility. The ‘cis’ arrangement of the DysF domains supports a dual membrane binding mode as previously proposed by Kaur et al based on in vitro lipid binding assays ^13^. In such a configuration, both DysF domains could dock onto the same membrane surface potentially increasing the retention of TECPR1 on damaged membranes. This could allow for multiple rounds of LC3 conjugation which is consistent with TECPR1’s role in CASM ^7–9^. Damaged endolysosomal membranes alter lipid packing and accessibility particularly SM exposure during CASM ^7,8,13,28^. The separation between the DysF lipid-binding sites suggests that TECPR1 may bind lipids with a fixed spacing. The distance between the predicted lipid binding sites is 98 Å (Figure 2B). This spacing allows TECPR1 to simultaneously engage at least two distinct sphingomyelin headgroups and could confer specificity towards the scale of membrane damage within the CASM pathway. Orthogonal studies have suggested that DysF1 is solely required for TECPR1 membrane recruitment. The close agreement between our DysF1 model and the previously solved crystal structure of TECPR1 DysF1 (PDB:8P5P) reported by Boyle et al., 2023 indicates that the fold of DysF1 is intrinsically stable and not affected by the assembly of full length TECPR1. In our structure, the DysF1 adopts a more extended conformation relative to TR1, whereas, DysF2 appears to collapse and is closely associated with the TR2 core (Figure 1C). This raises the possibility of a sequential docking model such that DysF1 mediates the initial membrane contact, followed by DysF2 engagement to stabilize TECPR1 on damaged membranes. In this sequential docking model, the DysF domains would need to be mobile, which is consistent with their lower resolution in our reconstruction.

To further investigate how TECPR1’s domain architecture supports its interaction with sphingomyelin-containing membranes. We performed molecular dynamics simulations in the presence of a lipid bilayer containing a ratio of lipid headgroups (7:2:1 POPC:POPE:DPSM). Using all-atom simulation, we observed minimal domain level changes across a 1µs trajectory (Figure 2C, Movie S2, S3). Overall, the architecture of TECPR1 remained stable with an RMSD <1.3 Å throughout the simulation. TECPR1 has a progressive diffusion across the lipid membrane and the DysF1 domain is observed detaching and returned to the membra surface. Notably, the DysF2 maintains contact throughout the trajectory (Figure 2C). The TR1-PH bridge remained intact throughout the trajectory with an RMSD at the interface <1 Å. Initially, we expected the PH domain to engage the membrane as previously proposed. However, TECPR1 rocked side to side but the PH domain remained unbound. We speculate that the elongated conformation may prevent the PH domain from docking. This would support past data that the PH domain does not engage the membrane in the absence of the ATG5-ATG12 conjugate^15^.Overall, the MD simulation support the model that full length TECPR1 directly associates with membranes through a cis DysF arrangement which is stabilized by the TR1-PH interaction.

### Future directions -

In conclusion, we present the first high-resolution structure of TECPR1 providing a foundation for understanding how the overall architecture of individual domains could influence LC3 conjugation. Our structure has revealed a novel TR1-PH interface that positions the PH domain far away from the confirmed lipid binding sites of the DysF domains. This model supports a cis arrangement of the lipid binding sites in the two DysF domains. It is intriguing to consider that this structure may represent an inhibited conformation of TECPR1 which is distinct from its role in binding ATG5-ATG12. The AIR binding motif (573-610) is connected to the start of the PH domain (611-717) (Figure 1a). In the elongated arrangement, it is hard to imagine how ATG5-ATG12 would gain access to lipid headgroups for LC3 conjugation. This is even true when TECPR1 rocks towards the membrane in the all-atom simulations. It is still unclear how binding of the ATG5-ATG12 conjugate will affect the positioning of TECPR1 domains, or how the PH domain modulates membrane interactions through potential conformational changes within TECPR1. The PH domain may undergo a conformational change to allow for membrane access while Dysferlin domains remain membrane anchored. Given that our structure is determined in the absence of sphingomyelin or a lipid bilayer, further investigation will be required to dissect this potential mechanism.

## Methods –

### Expression and purification of TECPR1

Human TECPR1 was expressed and purified from HEK293F cells as follows: pCAG plasmid encoding TECPR1 (amino acids 1-1,165) with a Twin-Strep-Flag (TSF) tag with a TEV cleavage site at the N-terminus was used for transfection. For a 1 L expression, 1000 µg of TSF_TECPR1 plasmid was mixed with 3000 µg of PEI and transfected into cells with density between 1.5 – 2 x 10^6^ cells/ml. Cells were grown for 48 hours on a shaker (160 rpm) at 37℃ with 5% CO_2_ media supplemented with 1% Antibiotic/Antimycotic and 1% FBS. Cells were harvested by centrifugation at 1000 g for 10 min, washed with 25 ml PBS. Cell pellet was lysed for 15 minutes in the cold in 25 ml buffer A (50 mM Tris-HCl, 150 mM NaCl, 5% glycerol, 1 mM TCEP, pH 7.4) supplemented with 1% Triton X and EDTA-free protease inhibitor tablet. Lysed cells were centrifuged at 15,000 rpm for 30 minutes, and the supernatant was added to 1 ml of streptactin resin and incubated for 1 hour in the cold by end-to-end rotation. The resin and protein mixture was transferred to a column and washed with 10 ml of buffer A. Elution was performed with 1 ml of buffer A supplemented with 10 mM D-desthiobiotin. Size exclusion chromatography was performed on a Sepharose 6 10/300 column equilibrated with buffer B (50 mM Tris-HCl, 50 mM NaCl, 5% glycerol, 1 mM TCEP, pH 7.4). Desired protein fractions were pooled and concentrated. Purified proteins were flash-frozen in liquid nitrogen and stored in -80 ℃ until use.

### CryoEM sample preparation and data acquisition

Prior to vitrification, 0.01 mM LMNG was added to 0.9 mg/ml of protein. Three microliters of the protein sample was applied to a glow-discharged gold grids (1.2/1.3, 300 mesh, UltrAuFoil) and plunge-frozen using the FEI Vitrobot mark IV at 4 ℃ and 100% humidity using a blot force of 10 and a blot time of 3 s. Imaging was performed on FEI Titan Krios G4 Cryo-TEM (ThermoFisher Scientific) operated at 300 kV, equipped with the Gatan K3 direct detector and a Gatan BioQuantum Imaging filter. Images were taken at a 30^°^ stage tilt. A total of 5,939 movies were collected at a pixel size of 0.97 Å and 130,000X magnification with a dose rate of 40 electrons/Å^2^ fractionated in 21 frames. The defocus ranged from -1 to -1.8 µm.

### CryoEM image processing

All data processing was performed with CryoSparc (v4.6.0) ^29^ installed on a custom GPU workstation. Briefly, dose weighting and constant transfer function CTF) estimation and micrograph curation were performed to remove images with poor CTF fit estimates and thick ice. Next, the blob picker algorithm was used to generate multiple 2D classes, and a few well-defined classes were used for template-based particle picking. Several rounds of 2D classifications were performed, and 650,000 particles were selected for the generation of three Ab-Initio 3D reconstructions. Of these three reconstructions, one showed an intact TECPR1 shape. The 3D reconstruction with 321,646 particles was subjected to multiple rounds of 2D classification. One class with 124,496 particles showed well aligned higher resolution particles and was non-uniformly refined to generate a global resolution of 3.26 Å based on the gold-standard Fourier Shell Correlation (FSC) at 0.143. The directional FSC (dFSC) plot shown in (Figure S2d) was generated on a remote webserver at (https://3dfsc.salk.edu). The density map was further put through the post-processing DeepEMhancer tool ^30^ to sharpen the map and reduce noise for model building and refinement.

### Model Building and Refinement

Initial domain structure was determined via AlphaFold2 (Uniprot ID: Q7Z6L1) ^31^. Model docking was performed using UCSF ChimeraX ^32^. Individual halves (TR1 1-385, TR2 605-1200) of TECPR1 were rigid body docked of using the “Fit in Map” option on ChimeraX. This process gave initial positions which were used in iterative modeling. DysF domains were left as is given the lower resolution of the electron density. TR1, TR2 and PH domains underwent several rounds of refinement first manually using Coot ^33^ followed by energy minimization using real space refinement on Phenix ^34^. Multisequence alignment was performed on sequences curated from NCBI Uniprot and aligned using ClustalO within the Seaview 5.1 software package. Figures were created using USCF Chimera X.

### Molecular simulations

The TECPR1-membrane system was prepared using the CHARMM-GUI Membrane Builder ^35^. The initial protein coordinates were positioned above a 7:2:1 POPC:POPE:DPSM lipid bilayer, and the system was solvated with TIP3P water molecules and ionized with 150 mM KCl. The full system contained ∼359,000 atoms, including 690 lipids, in a 150x150x168 Å^3^ unit cell. Energy minimization and equilibration were performed according to the CHARMM-GUI protocol. Following equilibration, a 1 microsecond production simulation was conducted in the NPT ensemble at 303.15 K and 1 atm with a timestep of 2 fs. NAMD3 ^36^ and the CHARMM36 force field ^37^ were used for all simulations.

## ABBREVIATIONS

AIR: ATG5 interaction region
CASM: Conjugation of ATG8s to Single Membranes
CryoEM: cryo electron microscopy
DysF: Dysferlin domain
PH: Plekstrin Homology domain
SM: Sphingomyelin
STIL: Sphingomyelin TECPR1–induced LC3 lipidation
TECPR1: Tectonin Beta-Propeller Repeat-containing 1

## Data Availability

The Cryo Electron Microscopy maps and structure model have been deposited in the EMDB-75438 and PDB-10SN, respectively. Protein expression constructs will be deposited to Addgene database. Raw electron micrographs are available upon request.

## Acknowledgements

We want to acknowledge the use of the Titan Krios G4 Cryo-TEM at the Electron Microscopy Core Facility, which is supported and administered by the Office of Research, Innovation, and Impact at the University of Missouri. This includes Dr. Min Su for technical support and training in microscopy. We want to thank Dr. Stephanie Gates, the members of the Yokom lab, and members of the Gates lab for helpful discussion and review manuscript figures.

