## SupplementalFigures for "Full length TECPR1 displays ‘cis’ Dysferlin domain architecture"

**Table S1: Cryo-EM Data collection and Structure Statistics**

| **Data Collection** | EMDB-75438 ,PDB-10SN |
|---|---|
| Microscope | Titan Krios G4 |
| Voltage (kV) | 300 |
| Total Dose (e–/Å$^2$) | 40 |
| Defocus Range (μm) | -1 to -1.8 |
| Pixel Size (Å) | 0.97 |
| **Image processing** | **Combined Data Sets** |
| Movies Collected | 13,121 |
| Micrographs Selected | 10,261 |
| Particles Extracted | 3,754,792 |
| Final Map Particles | 124,496 |
| Symmetry Imposed | C1 |
| FSC Threshold | 0.143 |
| Final Map Resolution (Å) | 3.26 |
| Resolution Range (Å) | 3.5 – 12.0 |
| **Validation** | |
| Ramachandran Favored (%) | 92 |
| Ramachandran Allowed (%) | 7 |
| Ramachandran Outliers (%) | 1 |
| Rotamer Outliers (%) | 2.5 |
| RMSD Bond lengths (Å) | 0.21 |
| RMSD Bond Angles (°) | 0.41 |
| Clashscore | 9 |

**Figure S1: Cryo-EM Data collection and Local Resolution**

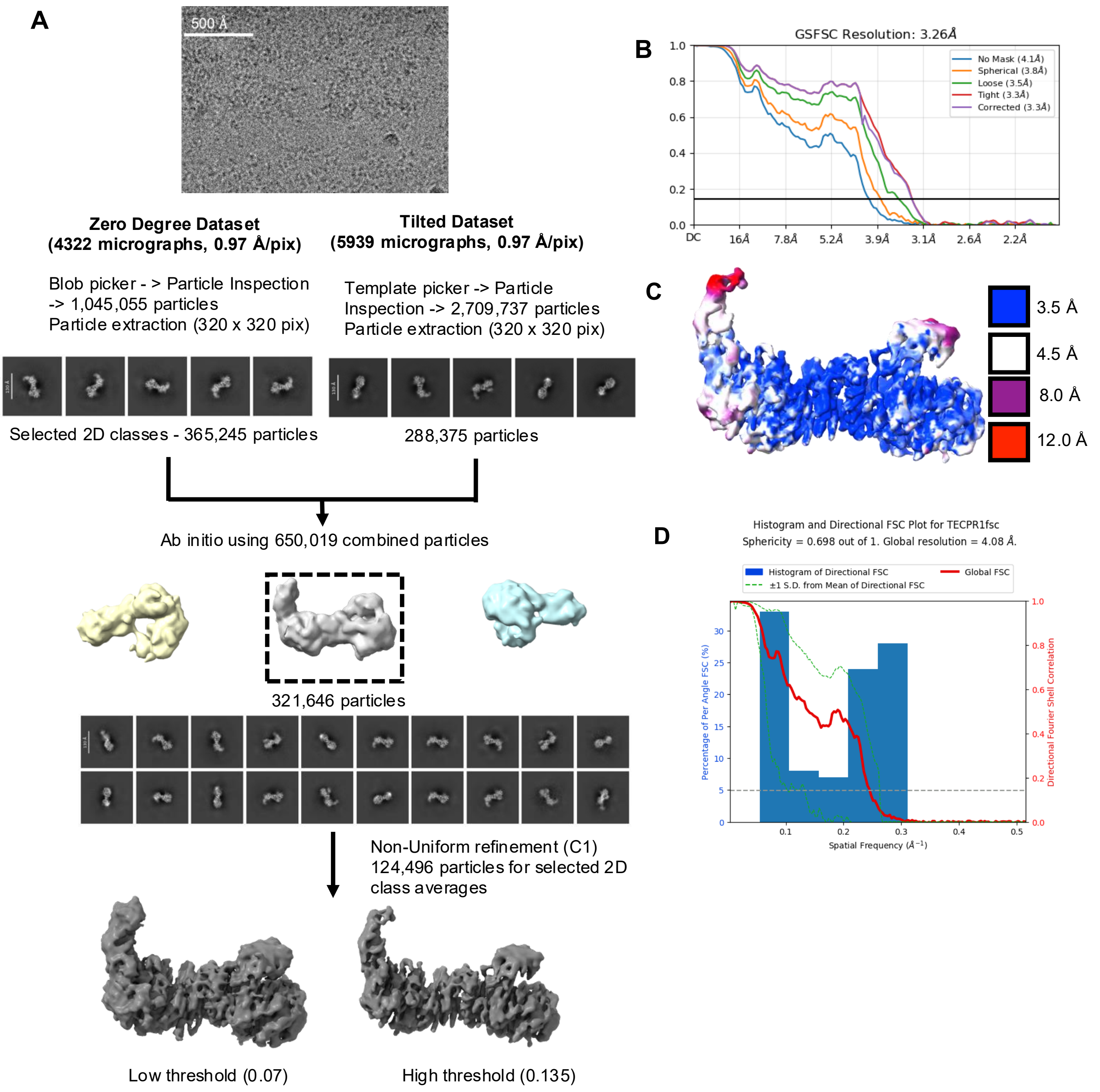

**Supplementary Figure S1. Data processing workflow of TECPR1.**

**A)** Representative cryo electron micrograph with TECPR1 protein. Data processing workflow used to generate the 3.26 Å map of TECPR1. 2D classes from the two datasets were combined, 365,245 particles from dataset1 and 288,375 particles from dataset2 were combined for a total of 653,522 particles. The 2D class averages were used for three (3) Ab-Initio maps generation. Selected Ab-Initio map was used for 3D refinements to obtain the final cryo EM map of TECPR1 containing 124,496 particles.

**B)** FSC curves calculated between independently refined half-maps under different masking conditions are shown: no mask (blue), spherical mask (orange), loose mask (green), tight mask (red), and mask corrected (purple). The final resolution of 3.26 was determined from the standard FSC plot with cutoff= 0.143, indicated by the horizontal black line.

**C)** Surface representation of TECPR1 density map coloured according to local resolution estimates. Regions resolved at higher resolution are shown at 3.5 Å (blue) and 4.5Å(white), while progressively lower resolution regions are indicated at 8.0 Å (magenta) and 12.0 Å (red).

**D)** Directional Fourier shell correlation (dFSC) analysis of TECPR1. Histogram (blue) shows the distribution of resolutions across Fourier space directions, while the global FSC curve (red) defines the overall resolution at FSC = 0.143. Dashed green lines indicates ±1 standard deviation of the directional FSC. The reconstruction has a global resolution of 4.08 Å and a sphericity of 0.698, indicating moderate directional anisotropy.

Supp Figure 2

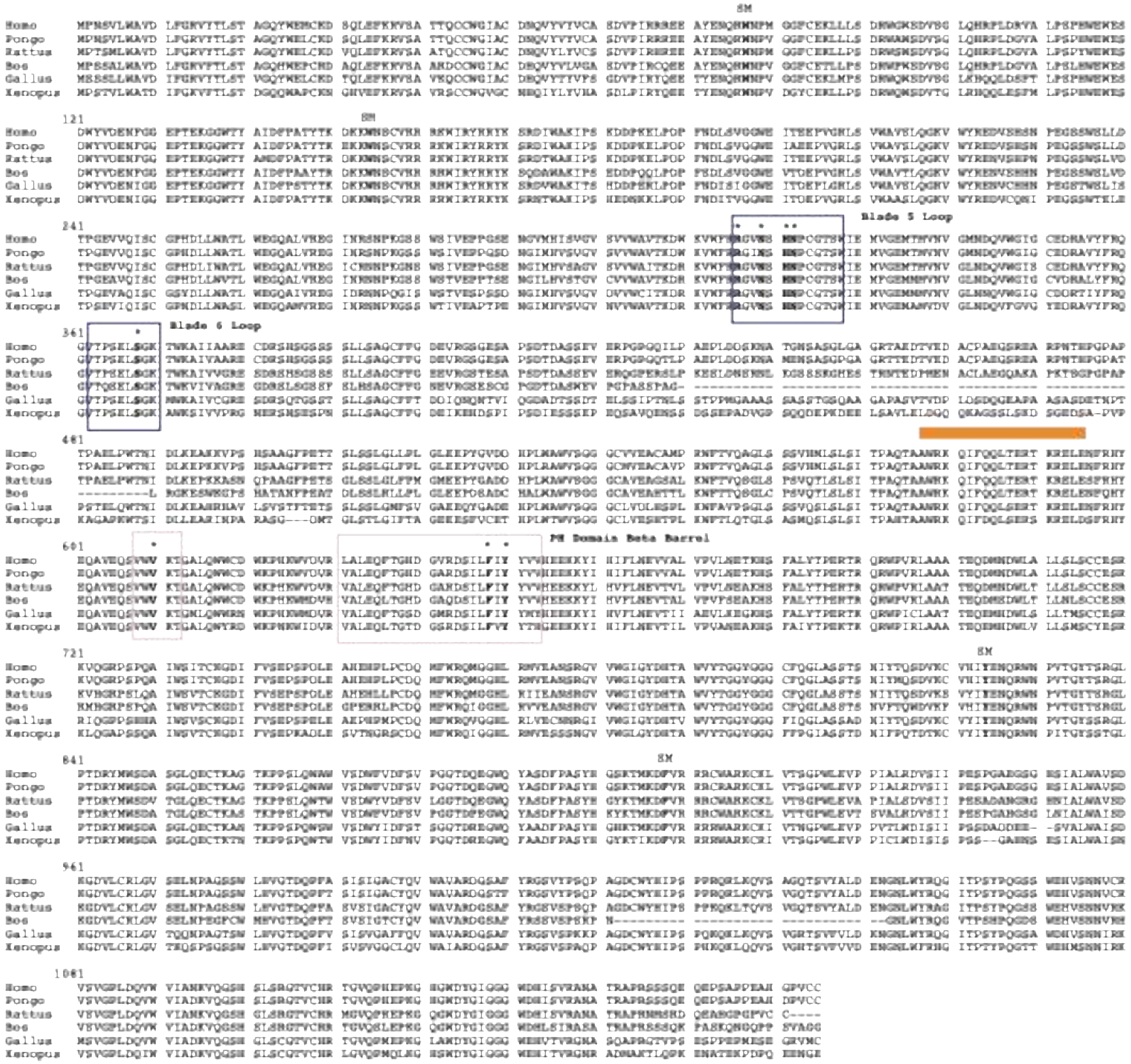

**Supplementary Figure S2. Sequence conservation of TECPR1**
Multiple sequence alignment of TECPR1 sequences from *Homo sapiens*, Rattus norvegicus, Gallus gallus demonstrates strong evolutionary conservation. Sphingomyelin residues are labelled with SM, TR1-PH interface residues are labelled with an asterisk, TR1-PH interfaces are colored corresponding to Figure 1A and AIR binding helix is shown with an orange box.

**Movie S1 – CryoEM density and model for full length TECPR1**

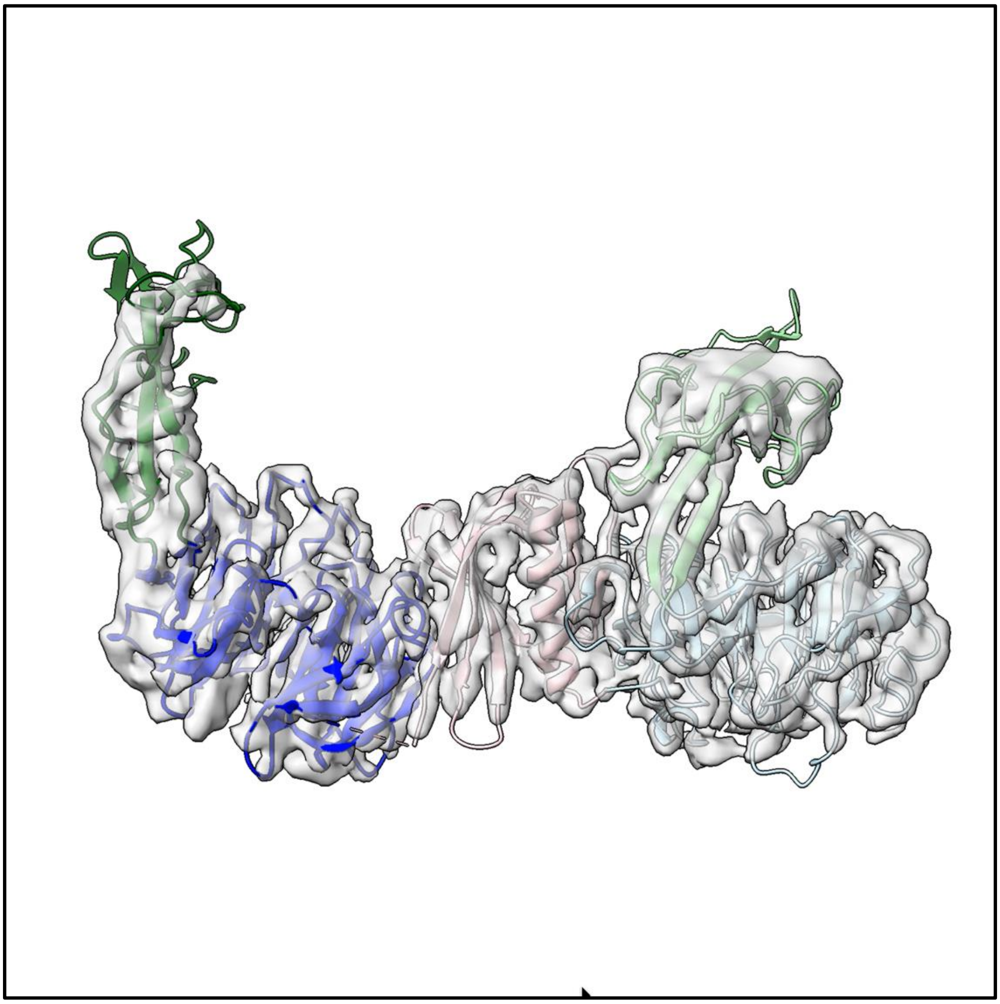

**Movie S2 – Side view of all atom simulation of TECPR1 at the membrane surface**

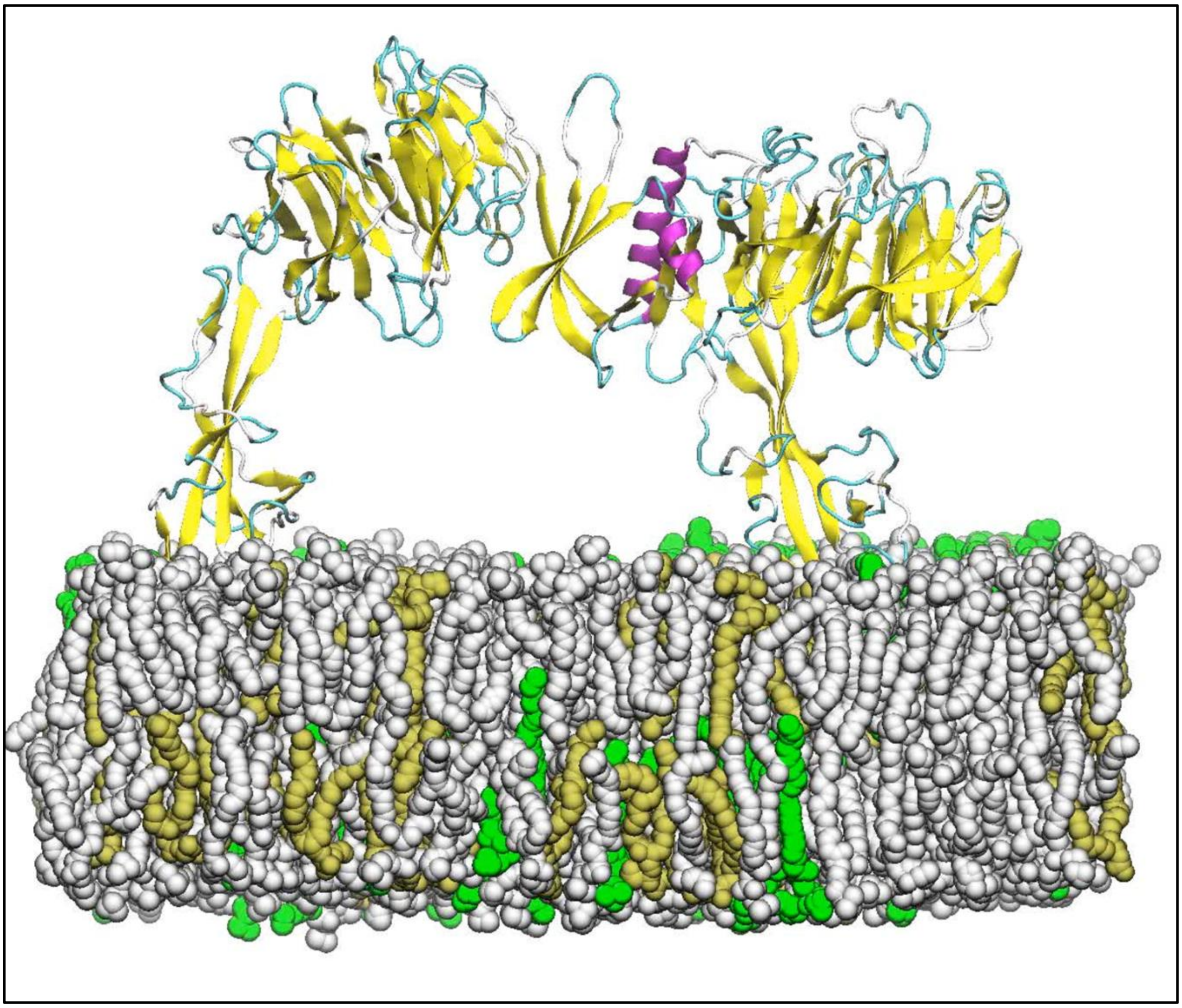

**Movie S3 – Top view of all atom simulation of TECPR1 at the membrane surface**

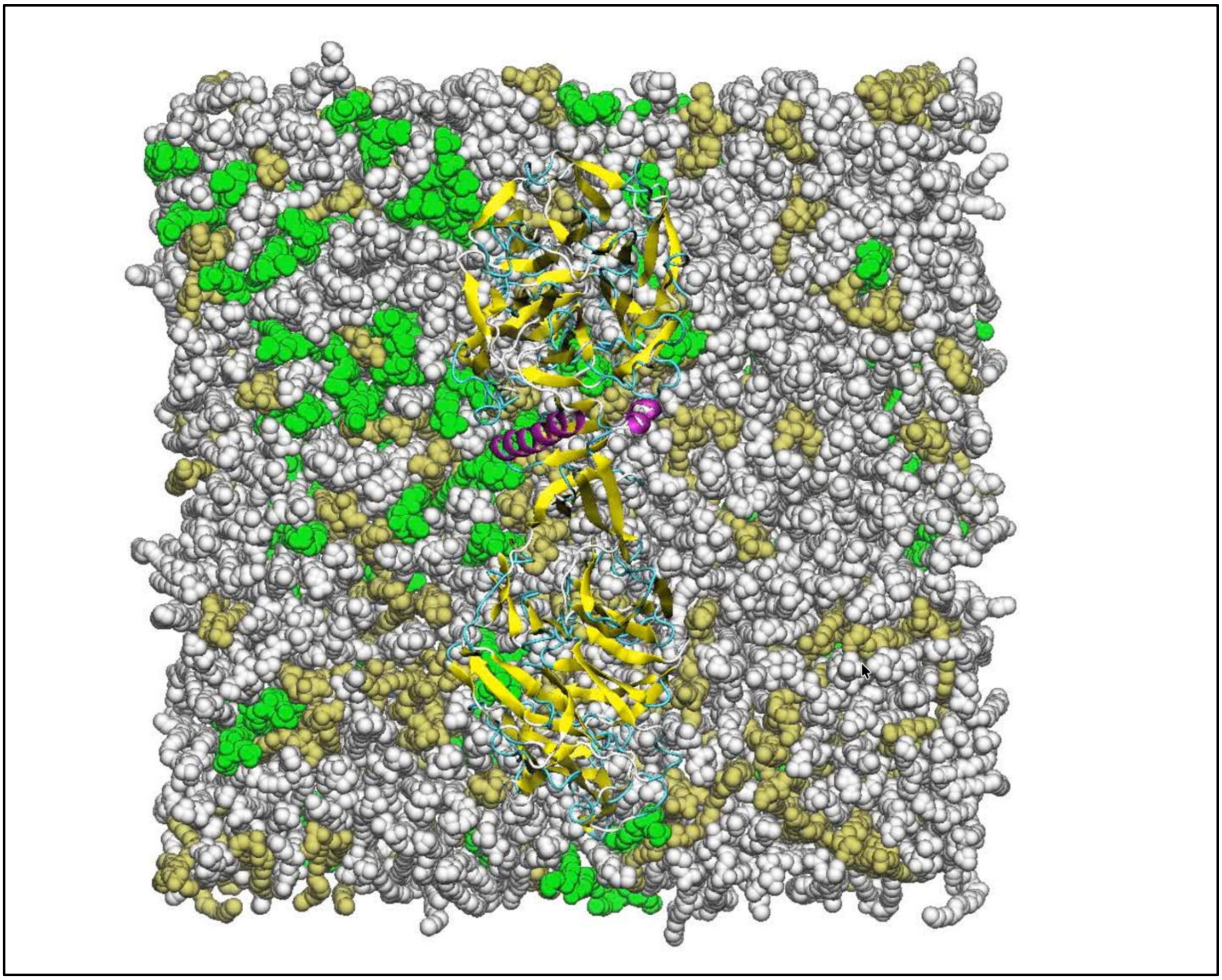
